# Measure Sodium Transport in Cells with NMR

**DOI:** 10.1101/2025.07.08.663659

**Authors:** Juntao Xia, Yu Yin, Yun Chen, Xuetao Wu, Haoyu Li, Yaqin Liu, Qingping He, Yun Ji, Ke Ma, Bona Dai, Hongzhen Bai, Fan Yang, Ruiliang Bai, Xueqian Kong

## Abstract

Sodium ions (Na^+^) are fundamental to numerous physiological functions, such as maintaining electrolyte balance, enabling nerve impulse transmission, and facilitating muscle contraction. Dysregulation of Na^+^ transport across cell membranes is implicated in a range of health issues, including metabolic syndromes, neurological conditions, and cardiovascular diseases. However, current methods for assessing cellular Na^+^ activity often face limitations; they can be invasive or fail to capture dynamic changes. In this study, we introduce a non-invasive ^23^Na nuclear magnetic resonance (NMR) methodology designed to directly quantify the transport rate of sodium ions in living cells. Our technique integrates relaxation exchange spectroscopy (REXSY) with a multi-site exchange model, enabling the investigation of Na^+^ transport dynamics on a timescale of sub-seconds. A key advantage is its ability to differentiate between intracellular and extracellular Na^+^ pools based on the endogenous NMR relaxation difference, thereby avoiding the need for potentially disruptive exogenous reagents. Experiments conducted on human cell lines successfully demonstrated the technique’s capacity to distinguish between various physiological states, such as when ion channels are pharmacologically blocked or activated. The resulting measurements of Na^+^ transport rates and intracellular Na^+^ fractions show a clear correlation with cellular metabolic activity, offering valuable quantitative markers for monitoring transmembrane ion dynamics *in vitro*.

## Introduction

Sodium ions (Na^+^) serve critical roles in living systems, underpinning vital processes like electrolyte homeostasis, the conduction of nerve potentials, muscle cell contraction, and broader cellular metabolism.^1–3^ These physiological actions are primarily orchestrated by sodium channels and associated transport proteins embedded within the cell membrane.^4^ Consequently, the malfunction of these channels, particularly voltage-gated sodium channels, can precipitate severe health conditions, including neurological disorders like epilepsy,^5,6^ cardiovascular diseases^7–9^ and chronic pain.^10^ Studies have shown that the abnormal expression of sodium channels may lead to cancer, Alzheimer’s and other diseases.^11–13^ Such pathological states often manifest as abnormal sodium ion activities, leading not only to imbalanced ion concentrations in different cellular compartments but also to unregulated Na^+^ transport across the cell membrane.^14,15^ Therefore, developing ion-specific analytical tools capable of measuring sodium transport is essential for understanding cellular functions and diagnosing ion channel-related pathologies.

While techniques such as patch-clamps, sodium-sensitive fluorescent probes, and ion-selective electrodes (ISEs) are commonly employed to study ion transport,^16–18^ they possess inherent limitations. These methods can suffer from low specificity, with signals often influenced by other ions like K^+^, Ca^2+^, or Cl?.^19^ Additionally, their invasive nature, requiring physical penetration or the introduction of non-native substances,^20,21^ restricts their applicability, particularly in complex biological systems or *in vitro* settings.

As an element-specific and non-invasive technique, ^23^Na nuclear magnetic resonance (NMR) offers a promising alternative for directly detecting sodium ions in biological systems.^22–24^ Indeed, ^23^Na magnetic resonance imaging (MRI) is already utilized in clinical research to investigate diseases linked to abnormal sodium metabolism.^25–27^ Despite this potential, a significant challenge within the MR community has been the effective differentiation of intracellular 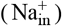 and extracellular 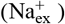 sodium pools, as these typically cannot be separated based solely on their intrinsic ^23^Na chemical shift (Fig. 1b). While parameters like non-intrinsic chemical shifts (using shift reagents),^28–31^ diffusivity^32^ and multiple-quantum transitions^33–36^ can, in principle, distinguish these pools, each comes with drawbacks. Shift reagents, such as lanthanide chelates, can have disruptive biological effects or interfering competitive ion effects.^37^ Diffusivity measurements using pulsed field gradients (PFG) are hampered by the rapid relaxation of intracellular Na^+^, rendering a significant portion undetectable.^38^ Multiple-quantum signal approaches, though capable of quantifying 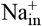, suffer from low signal-to-noise ratios and poor excitation efficiency.^39^

**Figure 1.**
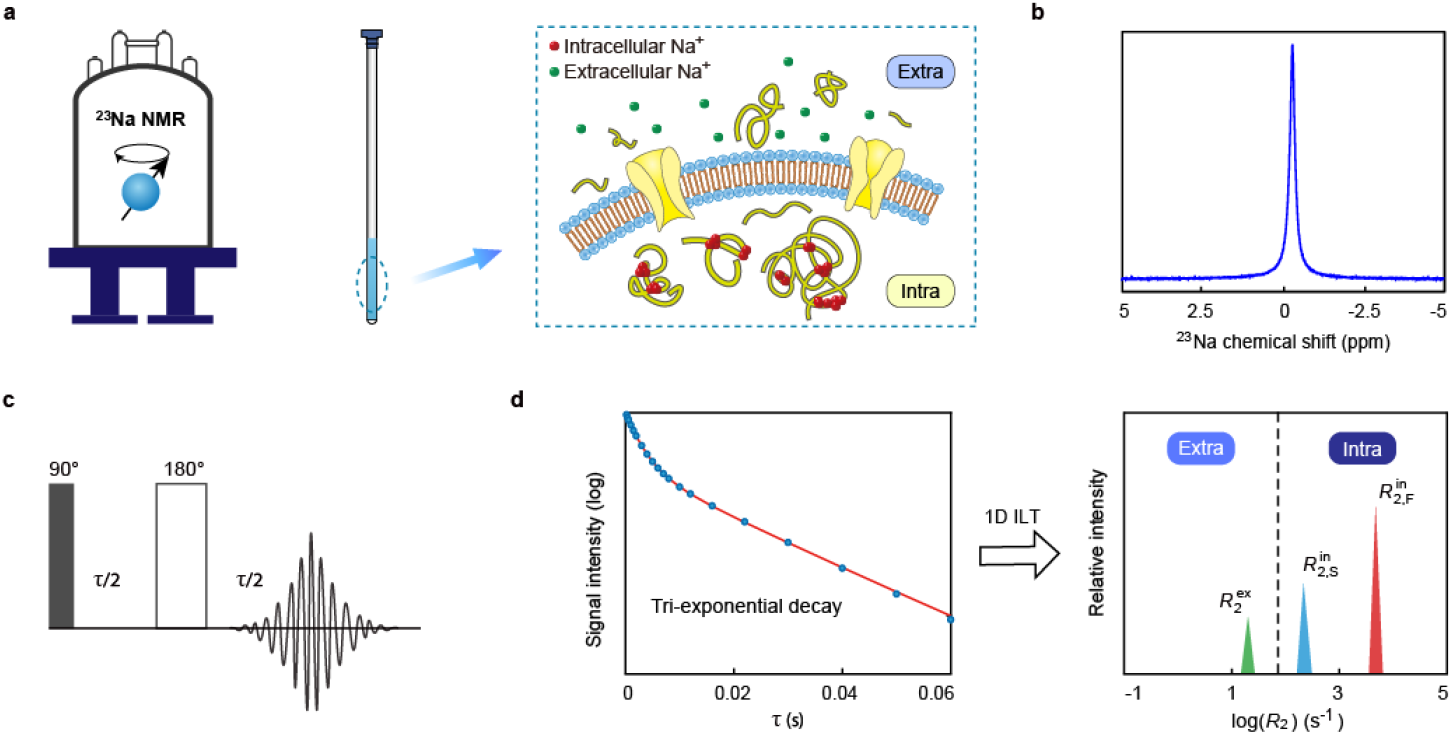
Schematics of ^23^Na NMR of living cells. (**a**) The NMR experimental setup and the illustration of sodium ions in the cellular environment. The red dots represent the intracellular Na^+^ associated with macromolecules while the green dots represent free extracellular Na^+^. The proteins on the cell membrane provide pathways for the transport of Na^+^. (**b**) ^23^Na NMR spectrum for sodium ions in living cells which shows a single resonance signal. (**c**) The spin-echo (SE) sequence with a variable delay time τ for acquiring the ^23^Na signal decay curve under *T*_2_ relaxation. (**d**) The fitting of the *R*_2_ decay curve either by a tri-exponential equation (the red curve in the left figure) or by the inverse Laplace transform (ILT) method (the plot on the right). The ^23^Na relaxation in a cellular system exhibits three *R*_2_ components, with two fast-decaying components that come from intracellular Na^+^ and one slow-decaying component that comes from extracellular Na^+^.

Our previous work established that the distinct multi-component relaxation behavior of ^23^Na spins serves as an effective parameter for differentiating 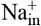 and 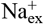. ^40–42^ The dynamic interactions between Na^+^ and intracellular macromolecules lead to relaxation properties markedly different from those of Na^+^ in the more dilute extracellular milieu. Building upon this, we have developed a time-programmable NMR pulse sequence that quantifies cellular Na^+^ dynamics using these relaxation parameters. By combining relaxation exchange spectroscopy (REXSY)^43–46^ with a multi-site exchange model, our approach directly probes Na^+^ transport rates on the millisecond-to-second timescale. Our findings demonstrate that the measured Na^+^ transport rates and associated parameters accurately reflect different cellular physiological states, such as those induced by ion channel blockers or activators. These quantitative metrics hold promise for *in vitro* monitoring of cellular activities. This study thus validates a non-invasive NMR strategy for sensing sodium activities in cellular assays, highlighting its potential utility in diverse areas of biomedical research.

## The Measurement Procedure

### The tri-exponential relaxation of cellular sodium

A standard NMR spectrometer (a 11.7 T solution-state NMR system with a 5 mm broadband probe) was used for the ^23^Na NMR measurement. Living cells were transferred from the culture dish into an NMR tube (Fig. 1a, see experimental section for details). A single frequency peak is observed in the ^23^Na NMR spectrum for the cell samples (Fig. 1b) as the intra- and extracellular Na^+^ could not be differentiated by chemical shift. Nevertheless, due to the stronger protein binding and higher viscosity inside cytoplasm, the intracellular sodium spins are influenced by quadrupolar interaction and it results with fast and bi-exponential transverse relaxation (*T*_2_).^38^ Our previous study showed that, for the cellular samples, the transverse relaxation of ^23^Na measured by the spin echo (SE) sequence (Fig. 1c) is sufficiently described by a tri-exponential decay function (Fig. 1d):

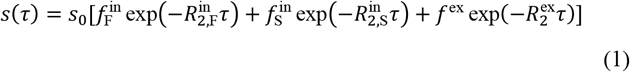

Here, *s*(τ) stands for the signal intensity at the echo time τ. Such tri-exponential decay also manifests as three well-separated peaks under inverse Laplace transform (ILT). The two relaxation rates 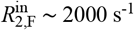 and 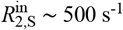 (with a fixed fraction ratio 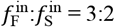, according to Redfield relaxation theory^47^) come from intracellular Na^+^; while 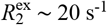 comes from extracellular Na^+^ in the culture medium.^38^ The fractions 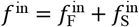 and *f*^ex^ represent the proportions of intra- and extracellular Na^+^, respectively.

### Measurement protocols of the relaxation-exchange sequence

During NMR experiments, the transmembrane transport of Na^+^ leads to an exchange from the slow relaxation rate to the fast relaxation rates, and *vice versa* (the process is illustrated in Fig. 2a). The exchange/transport rate can be measured by a relaxation-exchange pulse sequence (namely REXSY).^48^ In the REXSY sequence (Fig. 2b), a filter module (for a period τ_1_) is applied first, which suppresses the fast relaxation components (i.e. 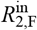 and 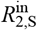) of intracellular signal; Secondly, an exchange module with a series of exchange periods *t*_m_ follows, which allows the partial recovery of fast relaxation components through the transmembrane transport; Finally, the detection module (for a period τ_2_) calibrates relative fractions of each relaxation component after exchange.

**Figure 2.**
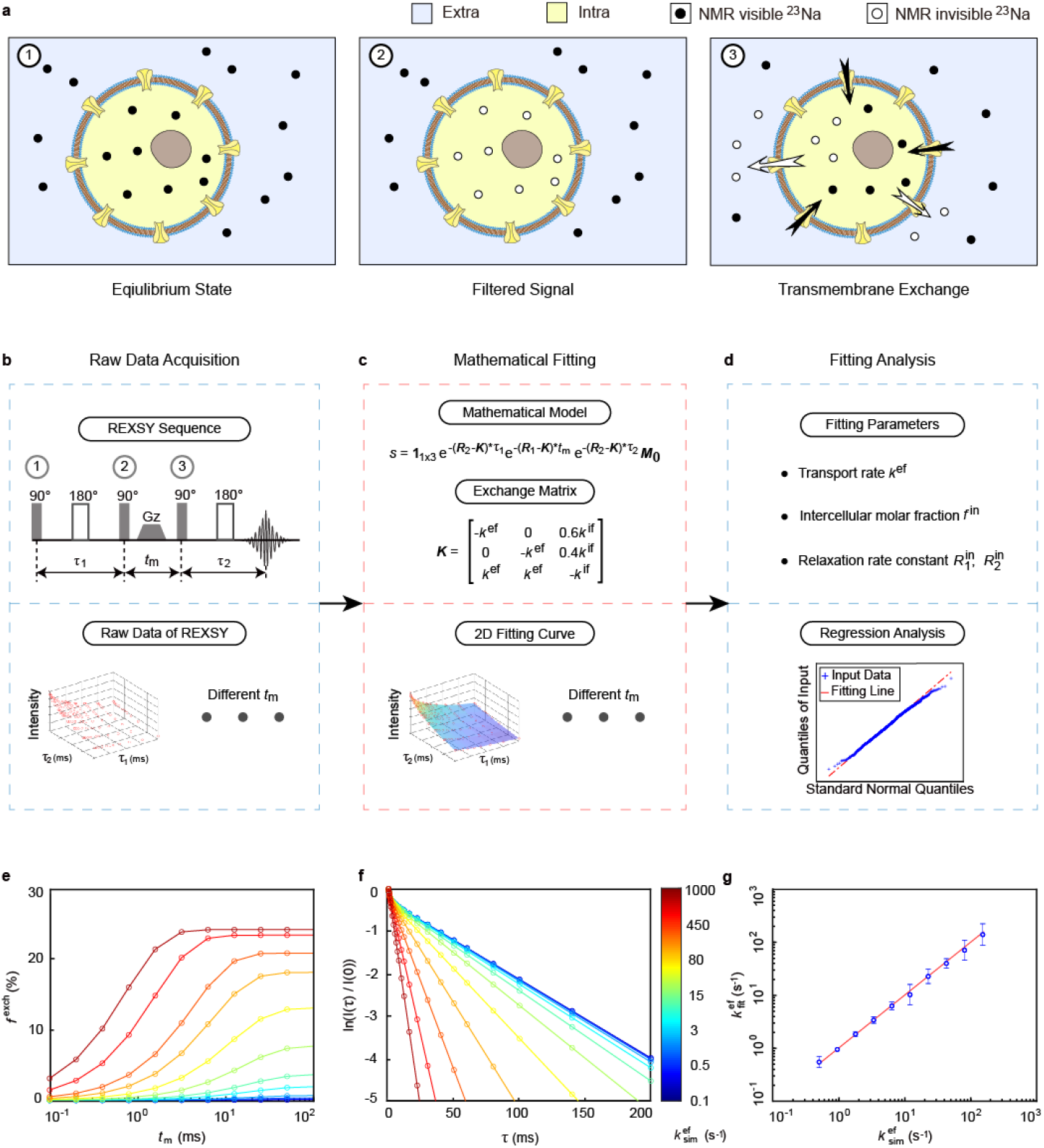
The measurement procedures of Na^+^ transport rate. (**a**) The illustration of cellular ^23^Na signal modulated by the REXSY pulse sequence shown in (**b**). The three stages correspond to the different modules of the REXSY pulse sequence. The NMR measurements are carried out in three steps: (**b**) signal acquisition, (**c**) curve fitting, and (**d**) data analysis. The fitting parameters are the transport rate *k*^ef^, intracellular Na^+^ molar fraction *f*^in^ and intracellular relaxation times 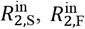 and 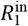. (**e**) The exchanged fractions and (**f**) the R^2^ decay curves for a range of simulated efflux transport rates 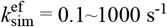. The following simulation parameters were used: 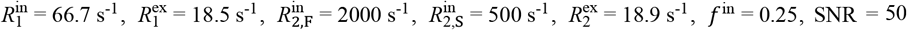. **(g)** The fitting values of 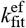 versus the simulated 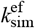 in the range of 0.5∼150 s^-1^. The results show good linear correspondence. The error bars come from the standard deviations of multiple simulation trials (n = 10).

Under REXSY measurement, the relaxation-exchange process can be described by the Bloch-McConnell equation:^49^

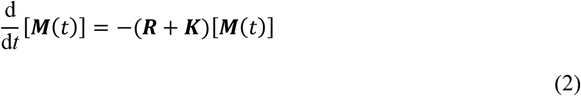

where ***R*** and ***K*** are the matrix representations of relaxation and transport rate constants. ***M***(*t*) is the magnetization vector at time *t*.

Here, we considered both longitudinal and transverse relaxation of ^23^Na in the relaxation-exchange process. The detected signal *s*(τ_1_, *t*_m_, τ_2_) is a three dimensional matrix as τ_1_, *t*_m_ and τ_2_ are varied during the measurement^44^ (Fig. 2c):

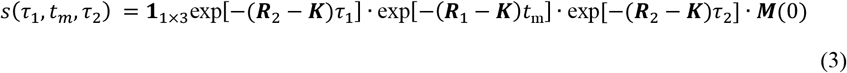

Here, **1**_1×3_ is a vector of ones.

In Eq. 3, the relaxation matrices are

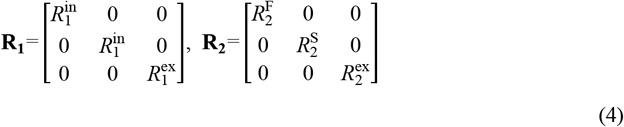

Intracellular ^23^Na has both fast and slow transverse relaxation rate constants (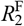 and 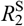, respectively), while has a single longitudinal relaxation rate constant 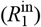. While extracellular ^23^Na has a single transverse relaxation rate constant 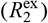 and a single longitudinal relaxation rate constant 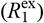.

The cross-membrane transport of Na^+^ is considered as the following dynamic equilibrium:

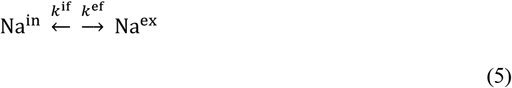

where *k*^ef^ and *k*^if^ are the Na^+^ transport rate constants for the cellular efflux and influx, respectively. Under the steady state, the rate constants satisfy:

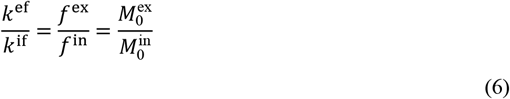

where 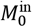 and 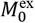 are initial magnetizations of intra- and extracellular ^23^Na, respectively, and they are linearly dependent on the Na^+^ molar fractions. In the following context, we name *k*^ef^ as the Na^+^ transport rate for simplicity.

Therefore, we can construct the exchange matrix and the initial magnetization vector:

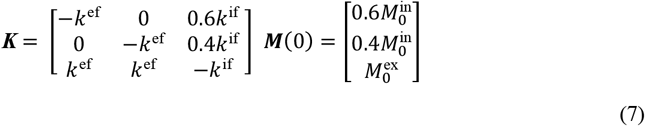

Note that the intracellular Na^+^ is divided into two fractions that satisfy the fixed ratio of fast and slow relaxation components:

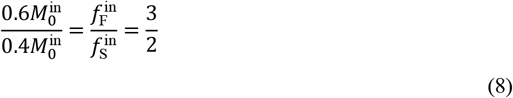

Upon setting up the measurement, we varied τ_1_ and τ_2_, both of which take values in the range of 0.1 ∼ 20 ms to obtain two-dimensional data at different *t*_m_ (2 ∼ 12 ms). The detected magnetization *S*(τ_1_, *t*_m_, τ_2_) was obtained as a 2D array and it was fitted by Eq. 3 using the regression fitting algorithm in MatLab. The 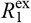 and 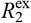 are predetermined by the inversion recovery and SE sequences: 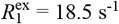 and 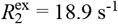. The rest parameters in Eq. 3 are fitting parameters. The quantile-quantile (Q-Q) plot (Fig. 2d) was used to assess the confidence and normality of fitting results.

To validate the efficacy of the REXSY measurement, we built a model that can simulate the REXSY process with arbitrary values of 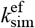. The model uses the relaxation parameters of typical HeLa cell samples. The lower bound of *k*^ef^ can be estimated in the plot of the intracellular exchanged fraction *f*^exch^ = *M*^in^*/*(*M*^in^ *+ M*^ex^) versus exchange period *t*_m_ (Fig. 2e). Here, *M*^in^ stands for the sum of the magnetizations of both 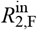 and 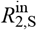 components, and *M*^ex^ stands for the magnetization of the 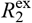 at the given time *t*_m_. For 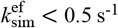, the exchanged fraction *f*^exch^ is hardly observable during a reasonable length of *t*_m_ (<20 ms). The upper bound of *k*^ef^ can be estimated by simulating the *R*^2^ decay (*I*(τ)*/I*(0)) of a spin echo sequence (Fig. 2f). Here, *I*(τ) stands for the magnetization at the echo time τ, and *I*(0) stands for the magnetization at τ = 0. The simulations show that, for 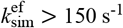, the *R*^2^ decay curve becomes a single exponential, which deviates from our tri-component relaxation model. Therefore, the REXSY measurement is applicable for 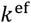 in the range of 0.5 ∼ 150 s^-1^. By applying fitting analysis to the simulation model, we obtain a good linear correspondence between the fitted value 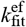 and the preset simulation value 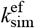 under a typical experimental condition with signal-to-noise ratio (SNR) = 50 (Fig. 2g).

### The cell cultures of different Na^+^ transport activities

In this work, we tested ^23^Na REXSY on human cervical cancer cell line HeLa and glioma cell line U-87 as well as the yeast cell line. The culturing procedures are given in the experimental section. In animal cells, the intracellular Na^+^ concentration [Na^+^] _in_≈ 10 ∼ 35 mM, while the extracellular concentration [Na^+^] _ex_ ≈ 140 mM. ^50,51^ For NMR experiments, we performed centrifugation to reduce the volume of culture medium and keep the fractions of intra- and extracellular Na^+^ at a comparable level. The procedure has minimal impact on the cell viability. The survival rate of the animal cells is over 75% even after the completion of NMR measurements (Fig. S6). For the HeLa cell line, we used three different conditions to activate or deactivate ion channels on the cell membrane:

- The treatment with steroid ouabain that partially blocks Na^+^/K^+^ ATPase to reduce the rate of Na^+^ transport (Fig. 4a).^52,53^
- The treatment with mannitol which is an agent that can induce hyperosmotic condition in cells and open the hypertonicity-induced cation channels (HICCs).^54^ In contrast, 2-aminoethoxydiphenyl borate (2-APB) can reverse the opening of HICCs (Fig. 4b).^55^
- The hypoxia treatment that induces cell swelling and eventually cell death.^56^ With the progression of the hypoxia condition, the cell membrane may be damaged leading to a significant increase of Na^+^ influx (Fig. 4c). Representative ^23^Na REXSY results and analyses are shown in Fig. 3a-3c and Fig. S1-S3. The Na^+^ transport rate *k*^ef^, intracellular Na^+^ relative molar fraction *f*^in^ and the intracellular slow transverse relaxation rate constant 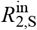 are compared for these treatments (see the Results session). The other fitting parameters are shown in the Supplementary Information (Fig. S4-S5).

**Figure 3.**
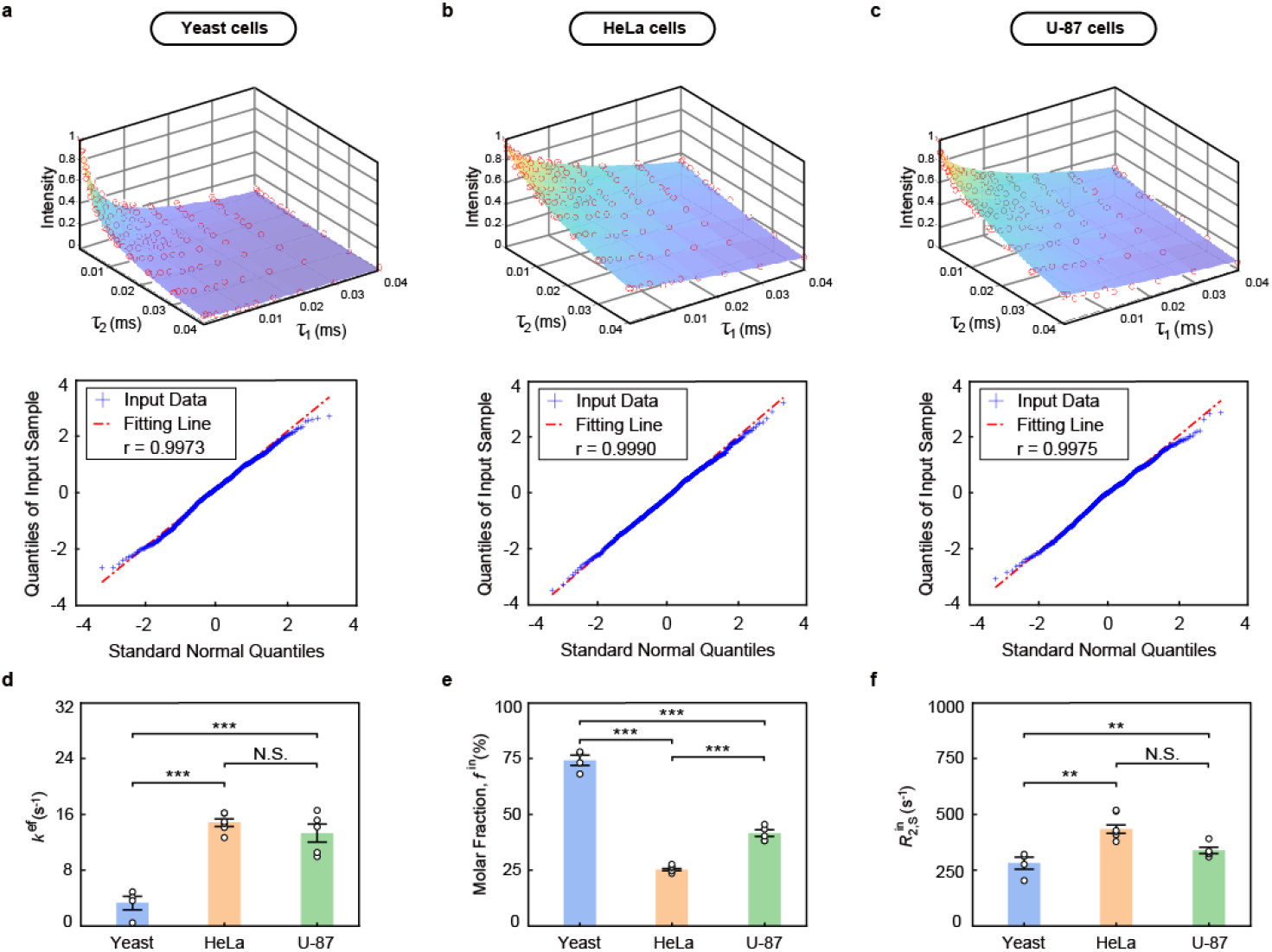
Sodium transport activities in different cell lines. The representative data sets of ^23^Na REXSY at *t*_m_ = 2 ms (above) and the corresponding Q-Q plots (below) for (**a**) yeast cells, (**b**) HeLa cells, and (**c**) U-87 cells, respectively. In Q-Q plots, data points closer to a linear distribution indicate better normality. (**d**) The transport rate constant *k*^ef^, (**e**) The molar fraction of intracellular Na^+^ (*f*^in^) and (**f**) The relaxation rate constants 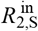 ^in^ in yeast group (n = 4), HeLa group (n = 5) and U-87 group (n = 5). The data were shown as mean ± s.e.m. * *P*<0.05, ** *P* <0.01, *** *P* <0.001. *P* values were calculated using one-way analysis of variance (ANOVA). N.S., not significant.

**Figure 4.**
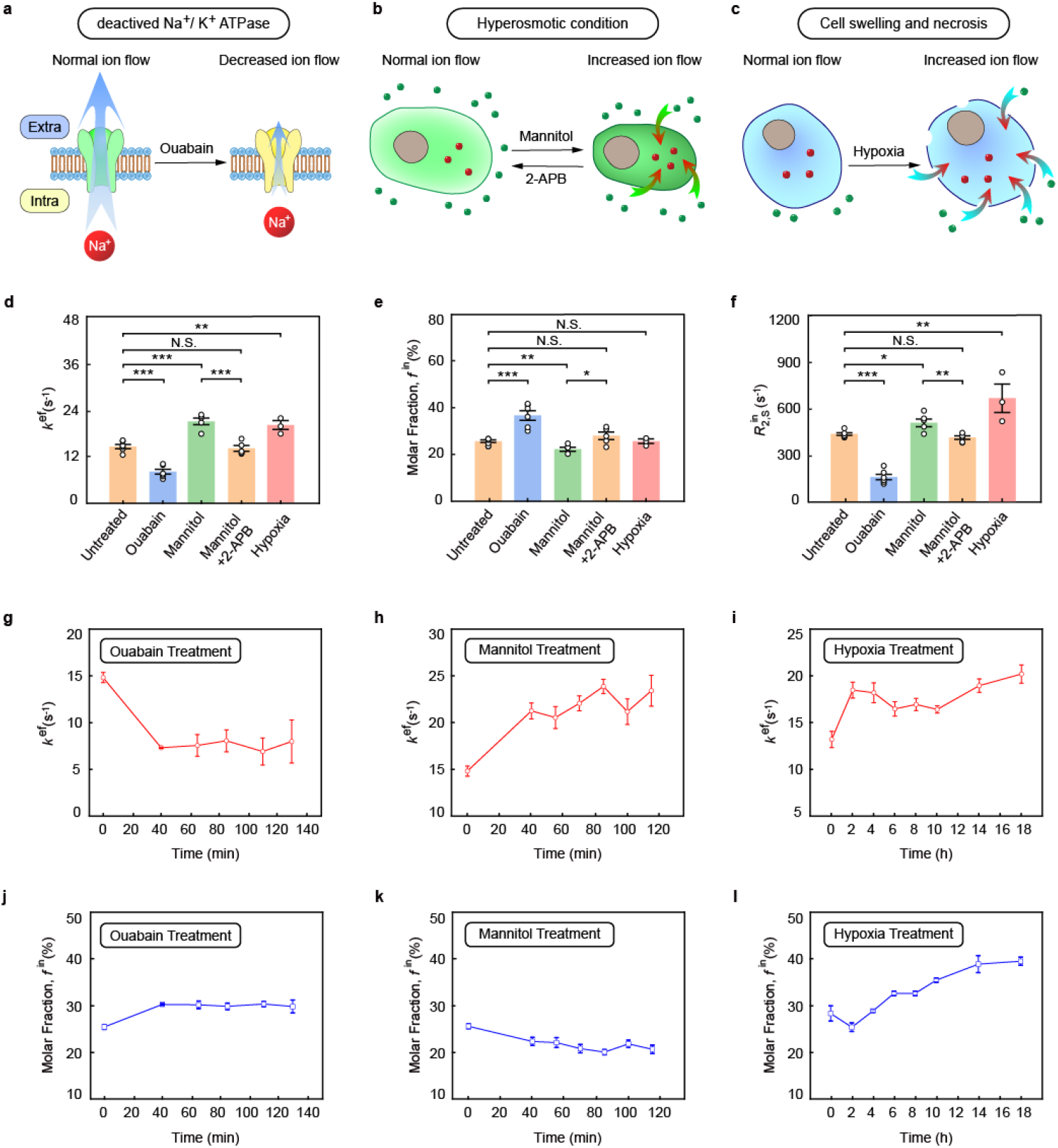
Na^+^ transport in cells under different treatment conditions. Schematic depictions of treatment conditions: (**a**) The treatment of ouabain which inhibits Na^+^ transport by blocking Na^+^-K^+^ pumps. (**b**) The hyperosmolar condition induced by mannitol and the reversal by 2-APB. (**c**) The necrotic process of cells under the hypoxic condition. The comparisons of sodium transport parameters: the bar charts of (**d**) the transport rate *k*^ef^, (**e**) the intracellular Na^+^ fraction *f*^in^ and (**f**) the relaxation rate 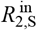 obtained with different treatment conditions: untreated group (n = 6), ouabain treatment (n = 6), mannitol treatment group (n = 5), mannitol+2-APB treatment group (n = 5) and hypoxia treatment group (n = 3). The time-lapse monitoring of Na^+^ transport activities: The changes of (**g**-**i**) the transport rate *k*^ef^ and of (**j**-**l**) the intracellular Na^+^ fraction *f*^in^ monitored during a period of time (n = 3 per condition). The data were shown as mean ± s.e.m. * *P*<0.05, ** *P* <0.01, *** *P* <0.001. *P* values were calculated using one-way analysis of variance (ANOVA). N.S., not significant.

## Results

### Na^+^ transport in living cells

Under the normal growing condition, the HeLa and U-87 cells have Na^+^ transport rates *k*^ef^ = 13 ∼ 15 s^-1^ (Fig. 3d) which are much higher than that in yeast cells (*k*^ef^ = 3.5 ± 1.1 s^-1^). The higher Na^+^ transport rates in HeLa and U-87 cells are associated with the elevated expression levels of sodium channels in mammalian cells (2 or 3 orders more abundant than those in yeast cells).^57–59^ We observe a high intracellular Na^+^ fraction in yeast cells (∼ 75%) as compared to HeLa cells (∼25%) and U87 cells (∼35%). The yeast cells were grown in a concentrated 1.8 M NaCl solution intentionally (Fig. 3e), and they cannot actively leach out intracellular Na^+^ since they lack of Na^+^ pumps.^60^ On the other hand, the abundant Na^+^ pumps on mammalian cells can actively regulate the Na^+^ balance between cytosol and extracellular environment.

The intracellular relaxation rate 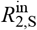 is a metric of bonding strength of Na^+^ within the intracellular environment.^38^ A higher 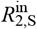 value indicates a stronger bonding of Na^+^ with proteins. The smaller 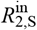 in the yeast cells (Fig. 3f) is likely due to the higher content of intracellular Na^+^, which leads to overall weaker bonding with intracellular proteins.^61^ 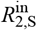 values in HeLa and U-87 cells are also different, which may indicate different cellular proteomes or different metabolic states for different cell lines.

### Na^+^ transport in cells with blocked ion channels

In the following experiments, we alter the condition of the HeLa cell line with different treatments, and test the ion activity with ^23^Na REXSY. When ouabain is used to deactivate the ion pump Na^+^/K^+^-ATPase, the Na^+^ transport rate in HeLa cells decreases to 8.1 ± 0.6 s^-1^ (Fig 4d). The remaining Na^+^ transport activity could be attributed to the contributions from other ion channels such as Na^+^, K^+^, 2Cl^-^-cotransport system^62^ or the incomplete blockage of Na^+^/K^+^ ATPase. Our result is consistent with the patch clamp study which showed the ion current decreased by 50% after adding ouabain to the frog bladder epithelial cells.^63^

The blockage of Na^+^/K^+^ ATPase leads to an increase of intracellular Na^+^ fraction (Fig. 4e), while the cell volume remains relatively constant as seen under microscope (Fig. S7). It means the intracellular Na^+^ concentration increases under the ouabain treatment. Meanwhile, 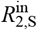 decreases from 438.9 ± 7.8 s^-1^ to 161.0 ± 17.6 s^-1^ (Fig. 4f) suggesting the bonding strength of Na^+^ ions inside HeLa cells becomes weaker.

During the course of NMR measurement (over 2 hours), the Na^+^ transport rate remains slow (Fig. 4g) and intracellular sodium molar fraction remains high (Fig. 4j). Our experiment confirms that ouabain has a lasting effect on the deactivation of Na^+^/K^+^ ATPase.^64^

### Na^+^ transport in cells with activated ion channels

As an opposite comparison, mannitol has been shown to induce the hyperosmotic condition which increases transmembrane ion transport and shrinks cell volume.^65,66 23^Na REXSY measurement shows that Na^+^ transport rate increases from 14.8 ± 0.5 s^-1^ to 21.3 ± 0.8 s^-1^. This is an indication of opened HICCs under the hyperosmotic condition and the state also manifests a substantially increased membrane current.^55,67^

Meanwhile, our measurement shows that intracellular Na^+^ fraction decreased by ∼ 20%, which may be attributed to the simultaneous activation of the cation-chloride cotransporters (e.g., NKCC).^65^ There is also an increase of 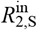, which indicates a stronger bonding of Na^+^ in the shrunken cells. We monitor HeLa cells under the hyperosmotic treatment for a period of 2 hours. The Na^+^ transport rate remains high (*>* 20 s^-1^) (Fig. 4h) and the intracellular Na^+^ fraction gradually decreases (Fig. 4k).^68^

In addition, studies showed that hyperosmotic condition can be reverted by the use of 2-APB drug.^55,67^ We show the effect of 2-APB can be also observed with the ^23^Na REXSY measurement. After the treatment of 2-APB, the Na^+^ transport rate returned to the normal state, so did the intracellular Na^+^ fraction and the 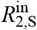 value (see Fig. 4d-4f).

### Na^+^ transport in cells under hypoxia

Hypoxia is a pathological condition that significantly impacts cellular homeostasis and functions.^69,70^ Our results revealed that hypoxia leads to notably different ion metabolism, as evidenced by a substantial increase (by 50%) of the Na^+^ transport rate (Fig. 4d). The rate remained at a high level since the initial stage of hypoxia treatment (Fig. 4i). The increased sodium activity can be attributed to the disfunction of Na^+^-K^+^ pumps under hypoxic necrosis conditions, which leads to higher membrane permeability of Na^+^ ions.^71–73^ This observation agrees with the electrophysiological study which showed an increase of membrane current after hypoxia.^74^

The intracellular Na^+^ first shows a slight decrease in the first 2-hour period and a later steady increase after 4 hours (Fig. 4l). The reduction in the first 2-hour period may be associated with the activation of the mitochondrial sodium-calcium exchanger (NCLX) during acute hypoxia.^75^ The increase in later period correlates to the onset of cell death, which is corroborated with the results of trypan blue staining (Fig. S6o). The influx of Na^+^ could be attributed to the cell necrosis process during which the cells swell and the membrane breaks down gradually.^56^ The process is also accompanied with an increased 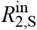 (Fig. 4f), which implies a dramatic change of proteins in the intracellular environment.^76–81^

## Discussion

Different from patch clamp and fluorescent techniques which mostly target single or a few cells, NMR is an ensemble measurement which targets more than 10^6^ cells simultaneously. Obviously, the Na^+^ transport rate measured by NMR is the average of all cells which neglects the heterogeneity in the sample. In another perspective, NMR can detect the collective activity of many cells rather than that of the very few selected one. The ion activity in cell clusters or organoids, which include a large quantity of internal cells inaccessible to microscope or micro-electrode, may be easily captured by the NMR technique.

The ^23^Na REXSY measurement is not just useful for the physiological study of cellular samples, but also applicable to many different biomedical systems. For instance, it can be used to probe the Na^+^ activities in extracted or engineered tissue samples. Furthermore, the REXSY sequence may be combined with MRI to map the Na^+^ transport rate or obtain Na^+^ transport weighted images in live bodies. Such REXSY-imaging (dubbed REXI) adaptation has been realized in the ^1^H detection to probe water transport in humans.^48^ The ^23^Na REXI protocol is currently under development.

In this work, the ^23^Na REXSY measurement of a single Na^+^ transport rate takes about 10 minutes. This measurement period accounts for the array of delay times that are needed for high-quality fittings as well as the repeated scans to increase SNR. It means that the measured Na^+^ transport rate is actually the average value over a period of ∼ 10 minutes. In principle, the measurement time can be reduced to less than 1 minute if the SNR is sufficient. The use of stronger magnetic field and better NMR probes (e.g. the cryoprobes) can improve the SNR. Using a larger sample volume (placing more cells into the NMR tube) also helps. On the other hand, the on-going development of CPMG-based sequence, which replaces the array of τ_2_ with a single scan, can further compress the measurement time down to ∼ 10 seconds. If fewer delay times are used, rapid measurement in the sub-second time scale could be achieved with a tradeoff of accuracy.

## Conclusion

In summary, the research established a versatile approach for calibrating Na^+^ activities in the cellular systems. The Na^+^ transport rate, the intracellular Na^+^ fraction and the intracellular Na^+^ bonding strength can be determined quantitatively. This approach can be adapted in different types of cells and reveal different metabolic states related to Na^+^ ions. This approach is ion-specific, non-invasive, and can be easily applied on a standard NMR instrument. In principle, this method can be combined with MRI to monitor the spatiotemporal Na^+^ activities in living organisms, which foresees great potential in medical research and diagnosis.

## Experimental Section

### Preparation of yeast cells

Commercial baker’s yeast (1.5 g, Angel Brand, China) was cultured in 1.8 M NaCl solution at room temperature for 4 hours. The intracellular Na^+^ concentration of yeast cells was measured to be 0.098 M using the previously report procedure.^38^ The yeast cell suspension was centrifuged at a speed of 3500 rpm for 3 ∼ 5 min. The supernatant in tube was decanted, and a total volume of 100 mL of 0.1 M NaCl solution was used to wash out remaining concentrated salt solution. After washing, the centrifugation was repeated. 300 mL of 0.1 M NaCl solution was added to the yeast cell precipitate to obtain the sample for NMR experiment.

### HeLa and U-87 cell culture

HeLa epithelial cervical cancer cell line and human U87 glioma cell line were obtained from the American Type Culture Collection (ATCC, USA). Both cell lines were cultured in 10 cm dishes containing 10 mL of high-glucose Dulbecco’s Modified Eagle Medium (DMEM, Meilunbio, China), supplemented with 10% fetal bovine serum (FBS, Sijiqing, China) and 1% penicillin-streptomycin (Solarbio, China). The Na^+^ concentration in the culture medium is 140 mM. The cultures were maintained at 37°C in a humidified atmosphere containing 5% CO^2^. The cells were routinely passaged every 3-4 days until reaching 80-90% confluence. For passaging, the spent medium was aspirated, and the cells were washed twice with 2 mL of phosphate-buffered saline (PBS, pH 7.4, Biosharp, China). Subsequently, 1 mL of trypsin (NCM Biotech, China) was added to the dish and incubated at room temperature for 30 seconds. After removal of trypsin, the dish was returned to the incubator for 2 minutes at 37°C to complete detachment. Digestion was terminated by adding 2 mL of serum-containing DMEM. The cell suspension was transferred to a centrifuge tube, pelleted at 1000 rpm for 5 minutes, and resuspended in 1 mL of fresh medium before reseeding. For NMR sample preparation, the cells were digested and centrifuged as described above. The pellet was resuspended in 200 μL of serum-containing medium and centrifuged at 1000 rpm for 1.5 minutes. The supernatant was carefully aspirated, and the cell pellet was transferred to a 5 mm NMR tube for analysis. All NMR experiments were conducted at room temperature without oxygen supplementation.

### Ouabain treatment

For the steady-state measurement, the HeLa cells were first washed twice with PBS and then cultured in 2.5×10^−5^ mM ouabain solution (purchased from Perfemiker, China; dissolved in serum-containing DMEM) for 3 hours at 37°C under 5% CO^2^. After that, the cells were digested and centrifuged, and then transferred into an NMR tube. For the time-lapse monitoring, the HeLa cells were first digested and centrifuged, and then soaked with 100 μL ouabain solution for a brief moment. After that, the cells were loaded in an NMR tube and resuspended with additional 200 μL ouabain solution. NMR measurements were started 40 minutes after adding the drug.

### Mannitol and 2-APB treatment

For mannitol treatment, the HeLa cells were first digested and centrifuged, and then the supernatant was replaced with 200 μL of 100 mM mannitol solution (prepared in serum-containing DMEM) for a brief moment. After that, the cells were resuspended in an additional 200 μL mannitol solution prior to NMR measurement.

For the 2-APB treatment, the HeLa cells were first digested and centrifuged, and then the supernatant was replaced with 200 μL of 100 mM mannitol solution for a brief moment. After that, the cells were resuspended in 100 μL of mannitol solution, incubated for 5 minutes, and then mixed with 100 μL of 2 mM 2-APB solution (in 100 mM mannitol solution) prior to NMR measurement.

### Hypoxia treatment

Following centrifugation, the pellets of HeLa cells were loaded into a 3-mm NMR tube. Supernatant was aspirated, and tubes were sealed with medical tape. A hypoxic environment was generated by evacuating air from the tubes using a syringe. Sealed 3-mm tube was then inserted into a 5-mm NMR tube which was further evacuated during experiments.

### Trypan blue staining

Cell viability was assessed post-experiment by mixing aliquots of the NMR tube suspension with trypan blue solution (BBI, Canada). Stained cells were counted under a microscope, and viability was calculated as the ratio of unstained (live) cells to total cells. HeLa cells in untreated, ouabain-treated, and hyperosmotic groups exhibited >90% viability after the completion of NMR experiments (Fig. S6).

### NMR experiments

The ^23^Na NMR experiment was carried out at 132.25 MHz using a Bruker AVANCE III system with an 11.74 T magnet using a broadband probe. The temperature was kept at 25°C during the experiments. At the beginning of the experiment, the 90° pulse was optimized (in the range of 10 ∼ 20 ms). The transverse relaxation time of ^23^Na was measured with the spin echo sequence using the echo times in the range of 0.1 ∼ 100 ms. For ^23^Na REXSY measurements, τ_1_and τ_2_ were set at different values between 0.1 ∼ 20 ms, while *t*_m_ was set at different values between 2 ∼ 6 ms. The spoiler gradient strength was 5×10^−4^ T/m. The recycle delay was set to 100 ∼ 150 ms. The number of scans was 16 or 32. Peak areas were obtained in TOPSPIN (Version 3.7.0, Bruker).

## Data analysis

All simulations and data processing were performed with MATLAB® (R2022a, the MathWorks, Natick, MA).

## Acknowledgements

This work was supported by the National Natural Science Foundation of China (22425402 and 22275159), Cyrus Tang Foundation (202523) and the Fundamental Research Funds for the Central Universities (YG2025ZD30).

