## SUPPLEMENTARY INFORMATION for "Measure Sodium Transport in Cells with NMR"

**Supplemental Figures**

**Supplemental Figure S1** The fitting results of ^23^Na REXSY measurement in HeLa cells with ouabain treatments.

**Supplemental Figure S2** The fitting results of ^23^Na REXSY measurement in HeLa cells with mannitol treatments and with 2-APB.

**Supplemental Figure S3** The fitting results of ^23^Na REXSY measurement in HeLa cells under the hypoxia condition.

**Supplemental Figure S4** The fitting parameters obtained from ^23^Na REXSY experiments.

**Supplemental Figure S5** Monitoring the changes of ^23^Na REXSY parameters.

**Supplemental Figure S6** Microscopic images of cells and the survival rates of HeLa cells under different treatments.

**Supplemental Figure S7** Changes in cell volume before and after ouabain treatment of HeLa cells.

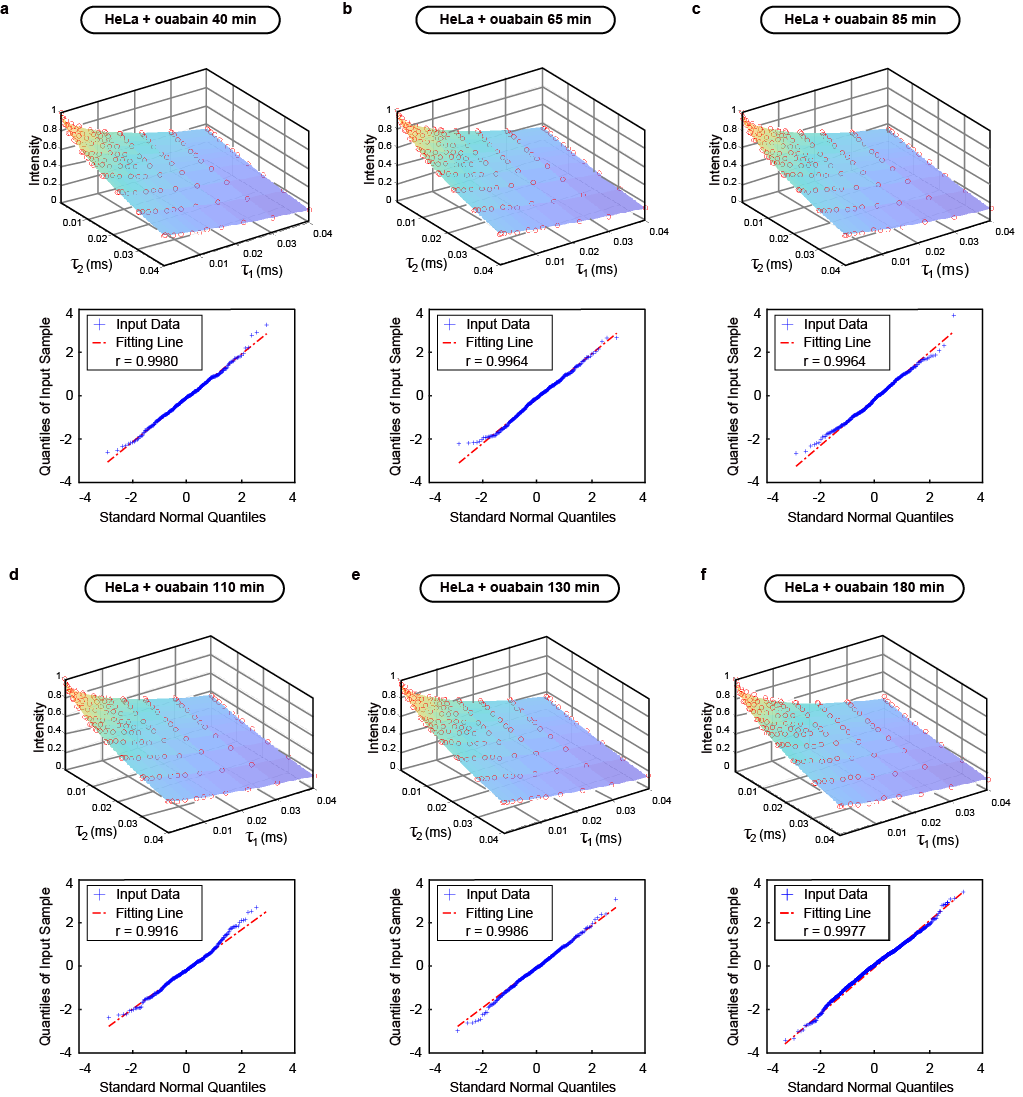

**Supplemental Figure S1** The fitting results of ^23^Na REXSY measurement in HeLa cells with ouabain treatments. A representative data set (at a selected $t_{\text{m}}\text{ = 2 }\text{ms}$, above) and the Q-Q plots (below) for ouabain-treated HeLa cells at (**a**) 40 min, (**b**) 65 min, (**c**) 85 min, (**d**) 110 min, (**e**) 130 min, and (**f**) 180 min, respectively. In Q-Q plots, data points closer to a linear distribution indicate better normality.

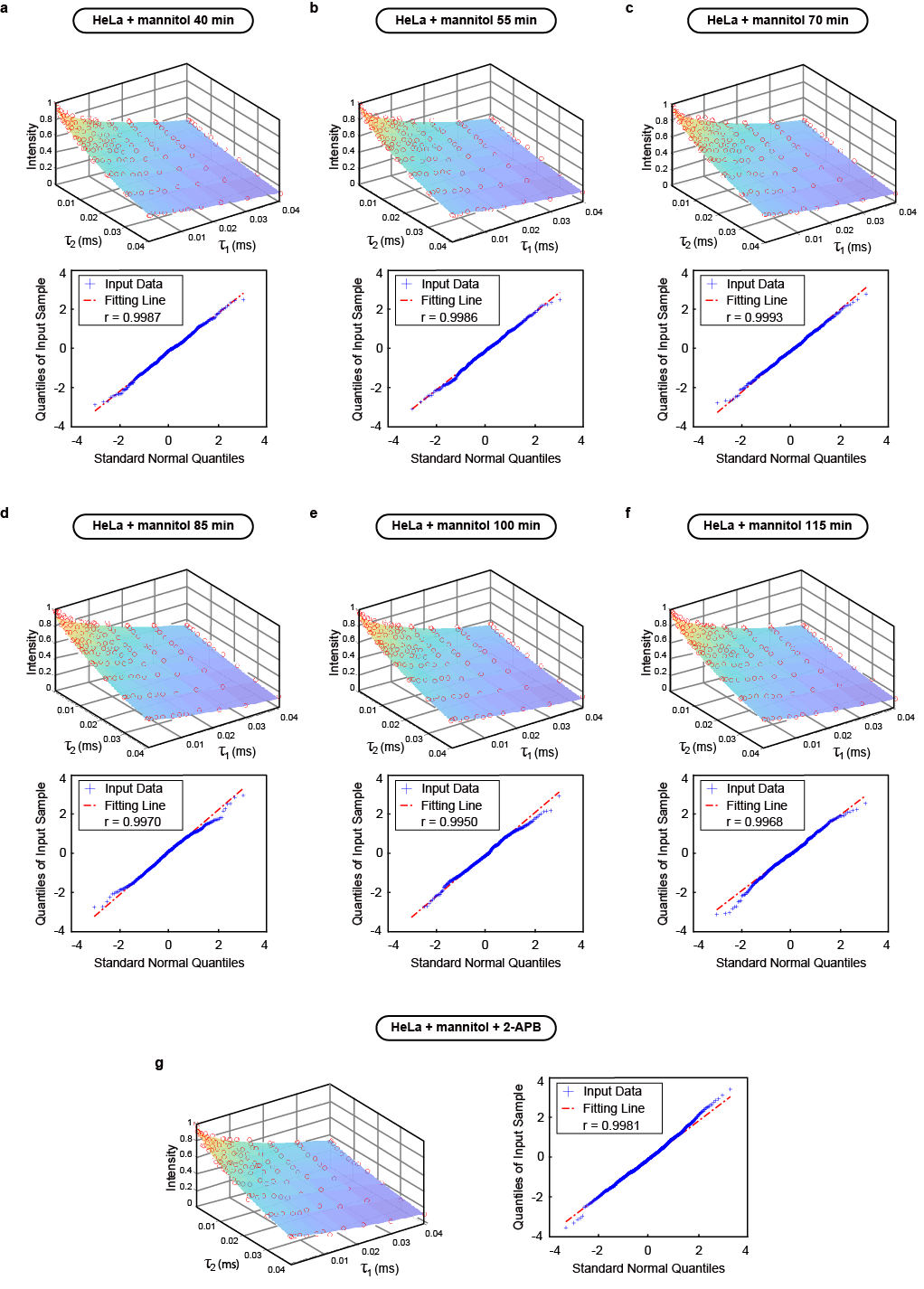

**Supplemental Figure S2** The fitting results of ^23^Na REXSY measurement in HeLa cells with mannitol treatments and with 2-APB. A representative data set (at a selected $t_{\text{m}}\text{ = 2 }\text{ms}$, above) and the Q-Q plots (below) for mannitol-treated HeLa cells at (**a**) 40 min, (**b**) 55 min, (**c**) 70 min, (**d**) 85 min, (**e**) 100 min, and (**f**) 115 min, respectively. In Q-Q plots, data points closer to a linear distribution indicate better normality. (**g**) the representative data set (at a selected $t_{\text{m}}\text{ = 2 }\text{ms}$, left) and the Q-Q plots (right) for mannitol-treated HeLa cells added 2-APB.

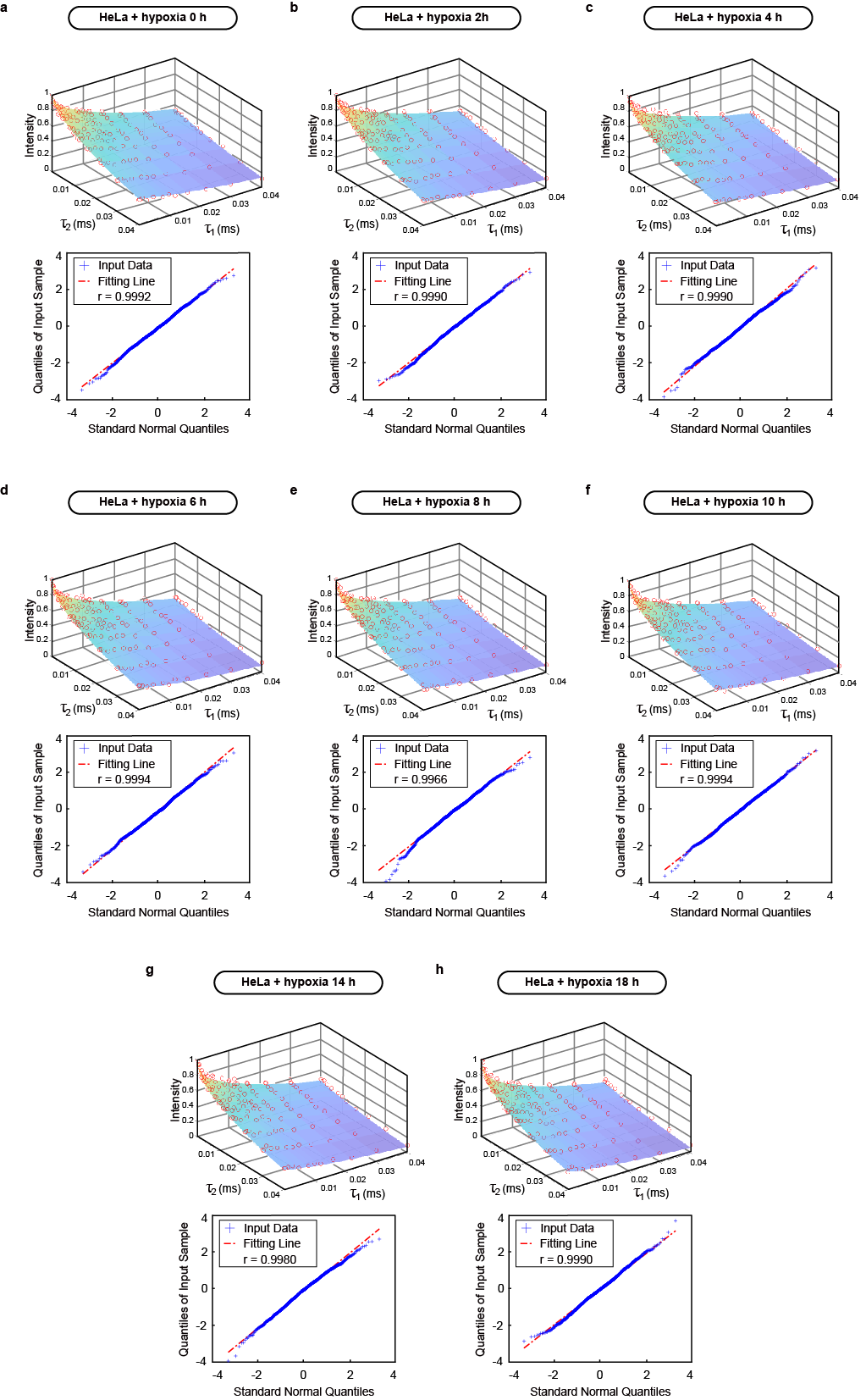

**Supplemental Figure S3** The fitting results of ^23^Na REXSY measurement in HeLa cells under the hypoxia condition. A representative data set (at a selected $t_{\text{m}}\text{ = 2 }\text{ms}$, above) and the Q-Q plots (below) for hypoxia-treated HeLa cells at (**a**) 0 hour, (**b**) 2 hours, (**c**) 4 hours, (**d**) 6 hours, (**e**) 8 hours, (**f**) 10 hours, (**g**) 14 hours and (**h**) 180 min, respectively. In Q-Q plots, data points closer to a linear distribution indicate better normality.

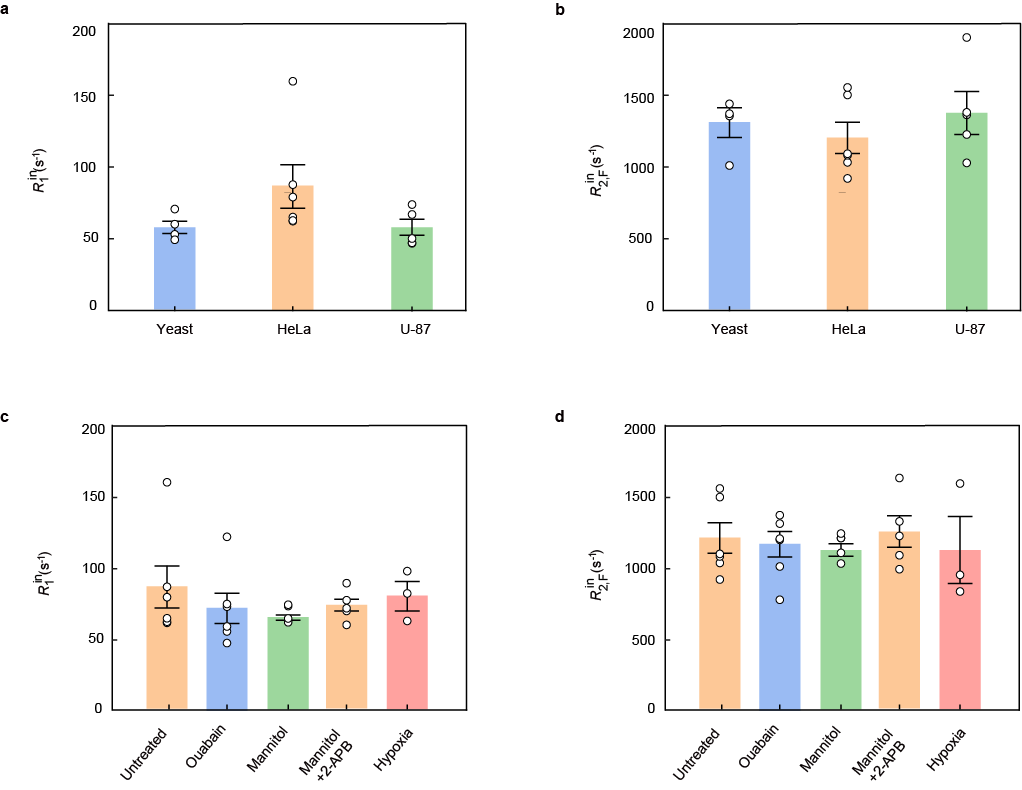

**Supplemental Figure S4** The fitting parameters obtained from ^23^Na REXSY experiments. The bar charts of (**a**) the longitudinal relaxation rate constant $\text{R}_{\text{1}}^{\text{in}}$ and (**b**) the transverse relaxation rate constant $\text{R}_{\text{2,F}}^{\text{in}}$ obtained from yeast group (n = 4), HeLa group (n = 5) and U-87 group (n = 5). The bar charts of (**c**) the longitudinal relaxation rate constant $\text{R}_{\text{1}}^{\text{in}}$ and (**d**) the transverse relaxation rate constant $\text{R}_{\text{2,F}}^{\text{in}}$ obtained in HeLa cells with different treatment conditions: untreated group (n = 6), ouabain treatment (n = 6), mannitol treatment group (n = 5), mannitol+2-APB treatment group (n = 5) and hypoxia treatment group (n = 3). The data were shown as mean$\text{± }$s.e.m. *P* values were calculated using one-way analysis of variance (ANOVA). All between-group comparisons showed no significant differences (*P* > 0.05).

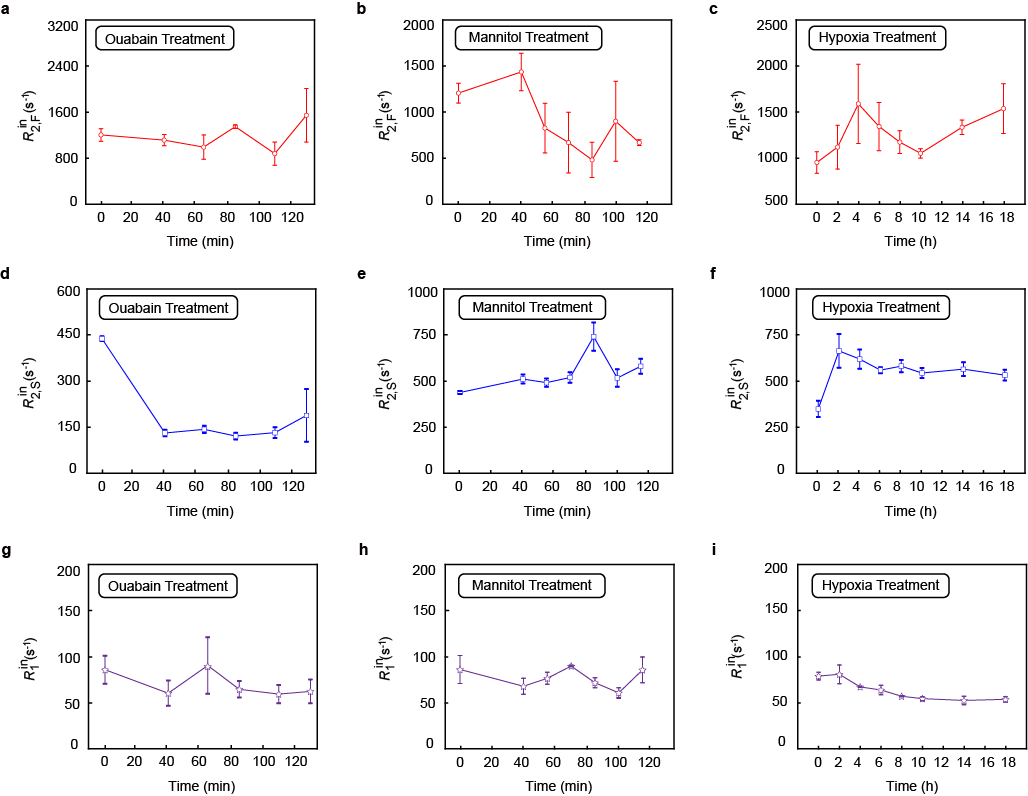

**Supplemental Figure S5** Monitoring the changes of ^23^Na REXSY parameters. (**a**-**c**) The fast intracellular transverse relaxation rate constants $\text{R}_{\text{2,F}}^{\text{in}}$, (**d**-**f**) the slow intracellular transverse relaxation rate constants $\text{R}_{\text{2,F}}^{\text{in}}$ and (**g**-**i**) the intracellular longitudinal relaxation rate constant $\text{R}_{\text{1}}^{\text{in}}$ for different treatment conditions (n = 3 per condition). The data were shown as mean$\text{± }$s.e.m.

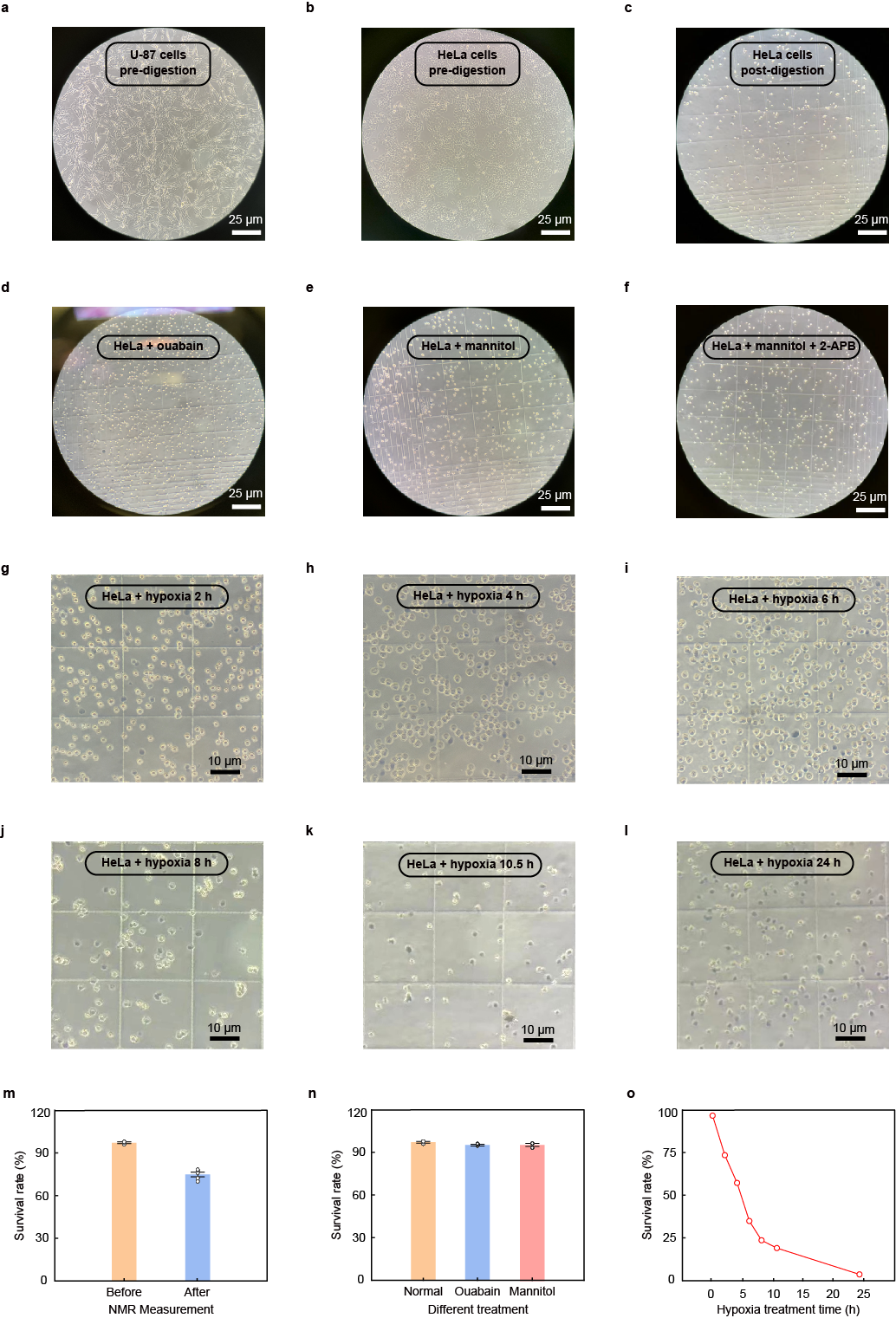

**Supplemental Figure S6** Microscopic images and survival rates of HeLa cells under different treatments. (**a**) Microscopic image of U-87 cells. (**b-f**) Microscopic images of HeLa cells under various conditions: (**b**) untreated group (pre-digestion), (**c**) untreated group (post-digestion), (**d**) ouabain-treated group, (**e**) hyperosmotic group, and (**f**) hyperosmotic group with 2-APB added. (**g**-**l**) Trypan Blue staining results at different treatment times. Cells stained with Trypan Blue indicate low viability, while unstained cells are viable. (**m**) Survival rates of HeLa cells before and after NMR measurements. (**n)** Survival rates of HeLa cells under different treatments. (**o**) Survival rates of HeLa cells under hypoxia during a period of time.

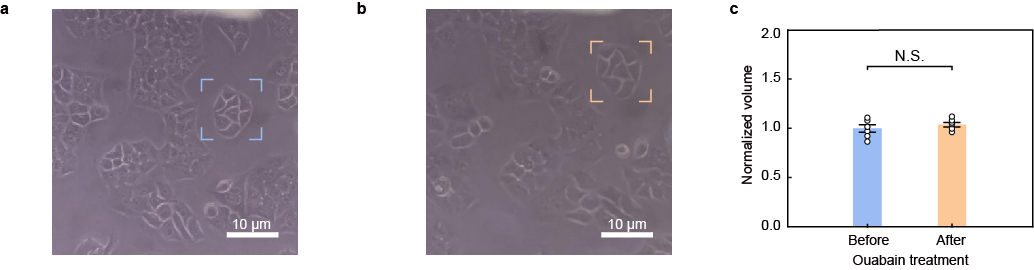

**Supplemental Figure S7** Changes in cell volume before and after ouabain treatment of HeLa cells. (**a**) The microscopic images of HeLa cells before ouabain treatment. (**b**) The microscopic images of HeLa cells after ouabain treatment. (**c**) the bar chart of cell volumes before and after ouabain treatment. The analysis shows that there is no significant change in cell volume before and after treatment. *P* values were calculated using one-way analysis of variance (ANOVA).
